# ACP-MHCNN: An Accurate Multi-Headed Deep-Convolutional Neural Network to Predict Anticancer peptides

**DOI:** 10.1101/2020.09.25.313668

**Authors:** Sajid Ahmed, Rafsanjani Muhammod, Sheikh Adilina, Zahid Hossain Khan, Swakkhar Shatabda, Abdollah Dehzangi

## Abstract

Although advancing the therapeutic alternatives for treating deadly cancers has gained much attention globally, still the primary methods such as chemotherapy have significant downsides and low specificity. Most recently, Anticancer peptides (ACPs) have emerged as a potential alternative to therapeutic alternatives with much fewer negative side-effects. However, the identification of ACPs through wet-lab experiments is expensive and time-consuming. Hence, computational methods have emerged as viable alternatives. During the past few years, several computational ACP identification techniques using hand-engineered features have been proposed to solve this problem. In this study, we propose a new multi headed deep convolutional neural network model called ACP-MHCNN, for extracting and combining discriminative features from different information sources in an interactive way. Our model extracts sequence, physicochemical, and evolutionary based features for ACP identification through simultaneous interaction with different numerical peptide representations while restraining parameter overhead. It is evident through rigorous experiments using cross-validation and independent-dataset that ACP-MHCNN outperforms other models for anticancer peptide identification by a substantial margin. ACP-MHCNN outperforms state-of-the-art model by 6.3%, 8.6%, 3.7%, 4.0%, and 0.20 in terms of accuracy, sensitivity, specificity, precision, and MCC respectively. ACP-MHCNN and its relevant codes and datasets are publicly available at: https://github.com/mrzResearchArena/Anticancer-Peptides-CNN.

## 1. Introduction

Cancer is one of the deadliest diseases in the world. Even though there are several ways of treating some of the cancer types, still there is no certain treatment for most of the cancers. Two of the major treatment strategies for cancer are radiation therapy and chemotherapy [1]. However, they are both expensive and have long term negative side effects [1]. In addition, cancer cells can become resistant to the chemotherapeutic drugs [1]. Therefore, there is a demand for finding new low cost and more effective treatments for cancer [2]. Among the newly introduced treatment methods for this deadly disease, anticancer peptides (ACP) have gained a lot of attention in the recent years as a less toxic and potentially more effective treatment for cancer [2, 3].

ACPs are short peptides consisting of 10 to 50 amino acids which are typically derived from antimicrobial peptides [4]. ACPs perform a wide range of cytotoxic activities against cancer cells while leave benign cells intact which is the reason behind their high specificity and low side effects [5]. Additionally, ACPs have low production cost, they are easy to synthesize and modify, and they have excellent tumour penetration capabilities [6]. In the past few years, many ACP based treatment options have been tested on a wide variety of cancer cells but only a few of them have been cleared for further clinical trials [7, 8]. Hence, rapid identification of potential ACPs is important for cancer therapeutic advancement. However, identification of these peptides through wet-lab experiments is relatively costly and time consuming [1]. Therefore, there is a demand for fast and accurate computational methods to tackle this problem. Among different computational methods, machine learning has merged as a promising approach to identify ACPs more efficiently and effectively.

During the past few years, a wide range of traditional Machine Learning (ML) methods have been proposed to identify ACPs. These traditional ML techniques require a set of hand-engineered features to represent protein sequences for the classification purpose. Thus, various methods for extracting effective features to represent proteins and peptides in an effective manner that contain significant discriminatory information for the classification purpose have been proposed. AntiCP was the first ML model for ACP identification that was proposed in [1]. In this model, peptide sequences are formulated by amino acid composition (AAC), split AAC (using N-terminal and C-terminal residues), dipeptide composition (DPC) and binary profiles features (BPF) [1]. Afterwards, these features are passed as input to a Support Vector Machine (SVM) classifier for separating the ACPs from the non-ACPs.

Shortly after that, Hajisharifi et al., proposed two methods for ACP identification using SVM [9]. In the first method, SVM was employed for separating ACPs from non-ACPs. They used pseudo-amino acid composition (PseAAC) method on different combinations of 6 physicochemical properties of the amino acids to extract their features. In the second method, the binary classification was performed using SVM with a local alignment based kernel method designed for feature extraction from peptide sequence. Later on, Chen et al. proposed iACP, where gapped dipeptide compositions (g-gap DPC) were used for feature extraction from peptide sequences, and SVM with radial basis function (RBF) kernel was used for the classification purpose [2].

More recently, Manavalan et al., proposed MLACP to tackle this problem. To build this model, AAC, DPC, atomic composition (ATC) of the sequences, and physicochemical properties of the residues were used for feature extraction while, SVM and Random Forest (RF) classifiers were used for ACP identification [10]. At the same time, Akbar et al., proposed iACP-GAEnsc, which used g-gap DPC, reduced amino acid alphabet composition (RAAAC), and PseAAC based on hydrophobicity and hydrophilicity of the amino acids (Am-PseAAC) for feature extraction. They also proposed an ensemble of different classifiers that combined SVM, RF, Probabilistic Neural Network (PNN), Generalized Regression Neural Network (GRNN), and K-nearest neighbour (KNN) classification models for ACP identification [11].

Later on, Xu et al., proposed a hybrid sequence-based model, where the peptides were converted to feature vectors through g-gap DPC to tackle this problem. They also used SVM and RF as their employed classifiers [12]. At the same time Kabir et al., proposed TargetACP, where the peptides were represented using split AAC, correlation factors extracted from PSSM profiles (PsePSSM), and composite protein sequence representation (CPSR). They also used SVM, RF and KNN classifiers as their employed models [13].

Most recently, Schaduangrat et al. proposed ACPred, where different combinations of AAC, DPC, PseAAC, Am-PseAAC, and physicochemical properties were used for peptide representation. They used SVM and RF classifiers for the ACP identification prediction [3]. At the same time, Wei et al., proposed ACPred-FL, where AAC, g-gap DPC, BPF, amino acid-specific physicochemical property-based bit vectors and composition-transition-distribution (CTD) methods were used for feature extraction. Similarly, they used SVM based ensemble model as their employed classifier [14].

Using traditional ML models (SVM, RF, KNN, etc.), the systems’ performances depend on the underlying manual feature extraction mechanisms. However, formulating problem-specific optimal feature representations for these sequences is not a trivial task and requires significant iterations of trial and error. In recent years, deep learning (DL) methods attracted tremendous attention to tackle challenging problems related to biological sequences because in many cases, unlike traditional ML algorithms, they do not require manual feature extraction to represent the input data [15–21]. Several DL methods, such as Convolutional Neural Network (CNN) [16, 22], Recurrent Neural Network (RNN) [16], word embedding [23, 24], and autoencoder [25, 26, 27] have been successfully employed for feature extraction and classification for DNA, RNA and protein sequences. Methods such as CNN and RNN exploit spatial locality and ordering information of the residues for ensuring that the extracted features retain a significant amount of discriminatory information from biological sequences.

However, none of the studies related to ML-based ACP identification explored automated feature extraction using DL methods until recently, when ACP-DL was proposed in [28]. To the best-of-our-knowledge ACP-DL is the only deep learning classifier proposed for this problem, so far. ACP-DL uses bidirectional long-short-term-memory (LSTM) recurrent layers for extracting features from peptide sequences followed by a fully-connected layer with a sigmoid neuron for classification. ACP-DL extracts features from two one-hot vector-based peptide representation techniques (binary profile and k-mer sparse matrix) that only depict the presence of a specific amino acid or a group of amino acids along different positions of the sequences. As a result, physicochemical properties or evolutionary substitution information of the residues, which contain significant signals regarding anticancer activities of peptide sequences are not utilized in ACP-DL’s feature extraction process [3, 14, 11, 13]. As a result, although the predictive performance of ACP-DL is quite impressive, there is still room for significant improvement.

Although recurrent layers are reliable for converting biological sequences into fixed-size features vectors [16], convolutional layers have also demonstrated promising performance addressing similar problems. In fact, CNN have been demonstrated as an effective technique for feature extraction and classification for DNA, RNA, peptides, and protein sequences in a wide range of studies [29–36]. However, CNN has never been used for ACP classification task. In this study, we hypothesize representation techniques that depict the residues’ evolutionary relationship and their physicochemical characteristics can embellish the feature extraction process for ACP identification since this type of information contains signals necessary for elucidating the structure and function of peptides. With this viewpoint, we are proposing a method called ACP-MHCNN, which consists of three jointly trained groups of stacked convolutional layers for interactive feature extraction from three distinct information sources for ACP identification. Our results demonstrate that ACP-MHCNN outperforms the current state-of-the-art methods on several well-established ACP identification datasets with a substantial margin. On ACP-500/ACP-164 benchmark dataset, ACP-MHCNN outperforms ACP-DL by 6.3%, 8.6%, 3.7%, 4.0%, and 0.20 in terms of accuracy, sensitivity, specificity, precision, and Matthews correlation coefficient (MCC), respectively. Our model and all its relevant codes and datasets are publicly available at: https://github.com/mrzResearchArena/Anticancer-Peptides-CNN.

## 2. Materials and Methods

In this section, we represent the benchmarks that are used in this study. We also present our sequence-representation methods as well as the proposed feature extraction and classification models.

### 2.1 Benchmark Datasets

In this study, we use three independent benchmarks to study the effectiveness and generality of our proposed method. These benchmarks are namely, ACP-740, ACP-240, and the combination of ACP-500 and ACP-164.

ACP-740 dataset was introduced in [28], consists of 740 samples out of which 376 are positive and 364 are negative. The positive samples (anticancer peptides) and negative samples (those without anticancer activity) in this benchmark are collected from [2, 25]. The ACP-240 dataset was introduced in [28], consists of 240 samples where 129 experimentally validated anticancer peptides are the positive samples and 111 AMPs without anticancer activity are the negative samples.

Two datasets, ACP-500 and ACP-164, were constructed in [14], where ACP-500 is used for training and validation, while ACP-164 is used as an independent test dataset. These two datasets consist of 332 positive and 1,023 negative samples, combined which are taken from [1, 2, 37]. Out of these samples, 250 positive samples and 250 negative samples are randomly selected for constructing ACP-500, whereas ACP-164 contains the remaining 82 positive samples along with 82 randomly selected negative samples.

### 2.2 Numerical Representation for Peptide Sequences

Although ACP-MHCNN does not require manual feature extraction, it is crucial to encode the sequences in numerical formats since the initial feature extraction layer of any DL architecture performs mathematical operations on the input for extracting class-discriminative activations. Such information is then passed as input to nodes in the subsequent layers. In this study, we exploit three peptide representation methods that are described in the following three sections. Since it has been shown in [14, 28] that considering *k* amino acids from the N-terminus of a peptide is sufficient for capturing its anticancer activity, we have represented each sequence using its *k* N-terminus residues. In our experiments, we have set *k* = 15.

#### 2.2.1 Binary Profile Feature (BPF) Representation

In our first representation method, each of the 20 amino acids (A, R, N, D, C, Q, E, G, H, I, L, K, M, F, P, S, T, W, Y, and V) is represented using a binary one-hot vector of length 20. For example, A is represented as [1, 0, …, 0], R is represented as [0, 1, …, 0], V is represented as [0, 0, …, 1], and so on. This representation encodes each sequence into a *k* × 20 matrix. Manually extracted short-range sequence patterns such as AAC, DPC, split AAC and long-range sequence patterns such as g-gap DPC have been successfully employed with traditional ML models for ACP identification [1, 2, 9, 12, 10, 14, 11]. We hypothesize that our model’s feature detection mechanism can capture both short-range and long-range sequence patterns that distinguish the peptides with anticancer activity from BPF representation.

#### 2.2.2 Physiochemical-based Representation

Basak et al., used a numerical representation for proteins for identifying length 5 conserved peptides through molecular evolutionary analysis [38]. The underlying numerical representation method proposed in [39] utilized an alphabet reduction strategy where the amino acids are divided into non-overlapping groups based on their side chain chemical property. The findings from these two studies have implied that amino acid physicochemical properties can facilitate the identification of evolutionarily conserved motifs, which are in turn important for maintaining the appropriate structure or function of the molecules. When these conserved motifs go through changes in the primary structure level, the amino acid residues are usually replaced with the ones with similar physicochemical properties. This phenomenon highlights the significant impact of exploring physicochemical properties for motif identification with respect to similarity among the substitute amino acids. Since our model identifies peptides with specific functions, discovering these motifs can strengthen our model.

Moreover, hand-engineered features based on amino acid physicochemical properties have been shown to improve ACP identification in a series of studies that have used traditional machine learning models [3, 9, 10, 14, 11]. We hypothesize that our feature extraction mechanism can identify similar features from a peptide representation based on the amino acids’ physicochemical properties. With these assumptions, our physicochemical property-based representation replaces each of the residues in a peptide sequence with a 31-dimensional vector (composed of 0/1 elements) that depict various physicochemical properties. As a result, each of the sequences is encoded into a *k* × 31 matrix.

For each amino acid, a unique 31-dimensional vector is formed through the concatenation of a 10-bit vector and a 21-bit vector. Elements of the 10-bit vector depict the membership of a specific amino acid in 10 overlapping groups based on its physicochemical properties as it was explained in [14]. Elements of the 21-bit vector are determined based on membership of a specific amino acid in the 7*3 = 21 groups formed by dividing them into 3 groups based on 7 physicochemical properties namely, polarity, normalized Van der Waals volume, hydrophobicity, secondary structures, solvent accessibility, charge, and polarizability as it was done in [14].

#### 2.2.3 Evolutionary Information based Representation

BLOSUM is a symmetric 20 × 20 matrix constructed by Henikoff et al., in [40], where each entry is proportional to the probability of substitution of a given amino acids with another amino acid in a protein (substitution probability in evolutionarily related proteins). Each entry in this matrix can be represented using the following equation:

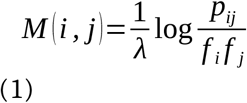

Where, *p*_*ij*_ is the probability of amino acids ‘i’ and ‘j’ being aligned in homologous sequence alignments, *f*_*i*_ is the probability that amino acid ‘i’ appears in any protein sequence, *f*_*j*_ is the probability that amino acid ‘j’ appears in any protein sequence, and *λ* is the scaling factor for rounding off the entries in the matrix to convenient integer values.

The observed substitution frequency for every possible amino acid pair (including identity pairs) is calculated from a large number of trusted pairwise alignments of homologous sequences as it is explained in [40]. If an entry M(i,j) is positive, the number of observed substitutions between amino acids i and j is more than random expectation. Thus, these substitutions are conservative (these substitutions occur more frequently than other random substitutions in homologous sequences). Therefore, each of the 20 rows of this matrix is a vector containing 20 elements that depict a specific amino acid’s evolutionary relationship with other amino acids. Here, we use BLOSUM matrix for retrieving a 20-dimensional vector for each of the 20 amino acids and use these vectors for encoding each peptide sequence into a *k* × 20 matrix. We hypothesize that our feature extraction architecture can automatically extract discriminative evolutionary features for ACP identification from this sequence representation. Among different BLOSUM matrix variations, we have used BLOSUM62 as the most popular one in this study.

### 2.3 Multi-Headed Convolutional Neural Network Architecture

CNN is a specialized neural network where each neuron in a given layer is connected to a group of neighbouring nodes in the previous layer. These layers drastically reduce parameter overhead and extract translation-invariant meaningful features by exploiting spatial locality structure in data through local connectivity and weight sharing [41]. A convolutional layer usually consists of several kernels where each kernel detects some specific local pattern in different input locations [41]. Since hand-engineered feature extraction methods such as AAC, DPC, g-gap DPC, PseAAC, and PsePSSM utilize ordering of neighboring residues and their correlation information with respect to evolutionary and physicochemical properties for feature generation from peptide sequences, using convolutional kernels for automatically approximating similar features is a rational choice. Moreover, well-defined ordering among the residues in peptide primary structure, the residues’ inherent local neighbourhood structures, and the presence of similar patterns (sequence motifs) at different locations across a peptide make these sequences perfect candidates for feature extraction through convolutional kernels.

The feature extraction mechanism in our proposed model consists of groups of stacked convolutional layers. Each convolutional layer group extracts features from a different representation of the peptide sequence. Since we have used three representation methods that serve as sources of discriminative information, our model contains three parallel layer groups. Each of these groups extract short-range and long-range patterns from a unique sequence representation using two stacked convolutional layers with varying number of kernels. The number of kernels in the layers and size of these filters are hyperparameters tuned through cross-validation [42].

The output feature maps of the second convolutional layer of each of the three groups are flattened, and the three resulting vectors are concatenated. The unified vector from this concatenation is passed through a dense layer with ReLU (Rectified Linear Unit) [43] activation function for recombining the features extracted from different sequence representations. It is to be mentioned that each element of the input vector for this dense recombination layer is calculated from a single information source (BPF or physicochemical or evolutionary representation) during forward-propagation. In contrast, elements of this layer’s output vector can be aggregated from multiple information sources. Hence, this layer enables seamless interaction between different convolutional groups that extract patterns from different representations and facilitates joint feature learning from multiple information sources during back-propagation [44]. These complex non-linear features are then passed as inputs to a dense layer with softmax activation function [45], which draws a linear decision boundary on the derived feature space for separating the anticancer peptides from peptides without anticancer activity. **Figure 1** represents the architecture of our proposed model for joint feature extraction from multiple information sources.

**Figure 1:**
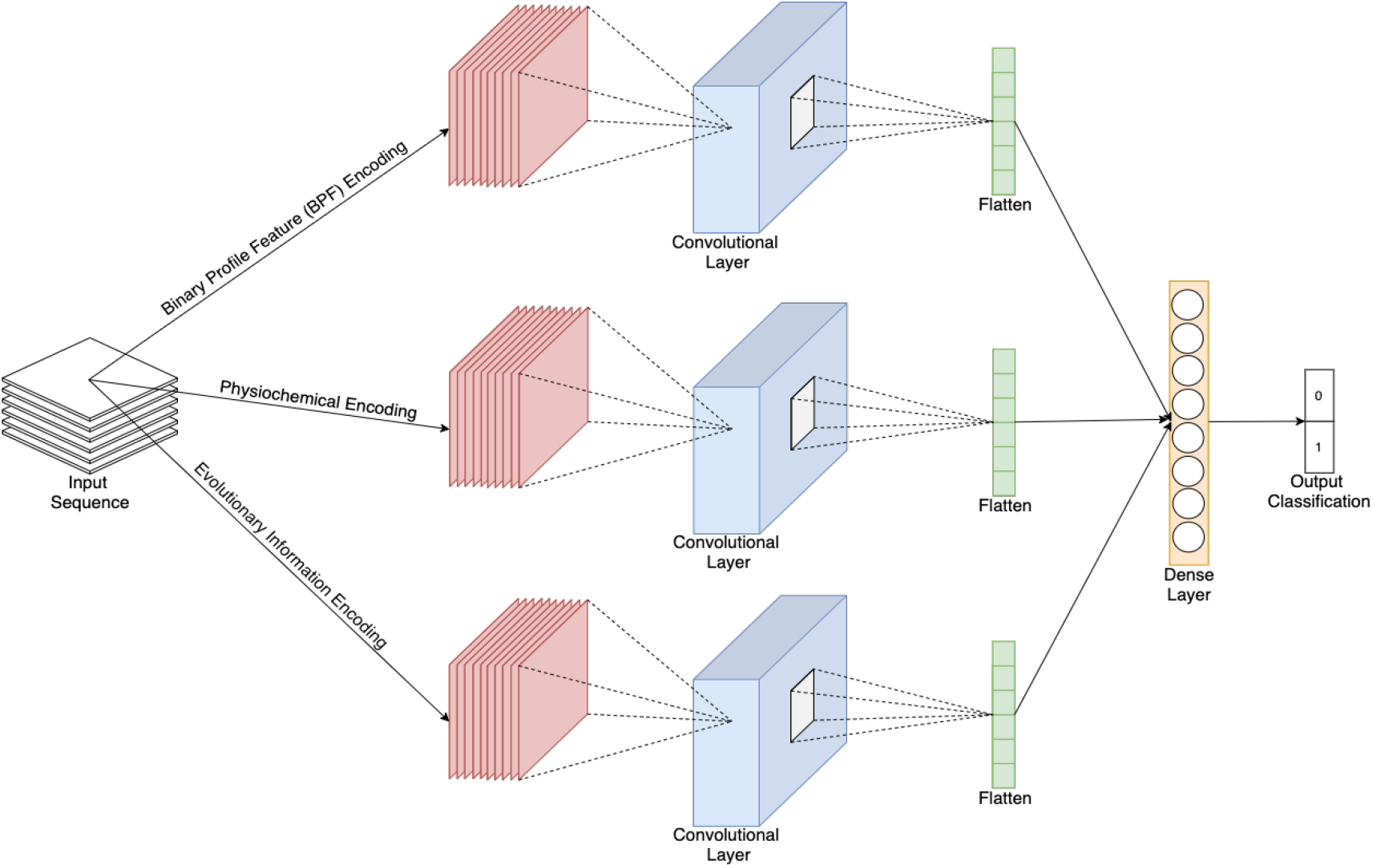
The general architecture of ACP-MHCNN. We extract BPF, physicochemical, and evolutionary-based features. We then feed the extracted features to a multi-headed deep convolutional neural network (MHCNN) to predict Anti-Cancer peptides.

Since the training data is limited for this task, there is a possibility for overfitting when training a deep-CNN model. To avoid overfitting, we adopt both L2 and dropout regularization methods in the feature extraction step to build out model [46]. L2 and dropout have been shown to be effective methods to address overfitting issue when the number of training samples are limited [46]. To be specific, the feature extraction occurs in all layers of the three parallel convolutional groups and the dense recombination layer after concatenation. Therefore, high dropout rates (>=0.5) are employed after each of these layers during the training phase to mitigate overfitting. These dropout rates are determined through cross-validation. Note that, the three convolutional layer groups that extract features from three distinct sequence representations are jointly trained alongside the dense recombination layer for minimizing cross entropy loss function [47]. Therefore, our model can simultaneously interact with the three information sources for detecting complex and ambiguous patterns. Optimal values for our model’s weights and biases are learned by employing Adam optimizer [44] with a learning rate determined through cross-validation.

ACP-DL, the only deep learning-based architecture proposed to date for anticancer peptide identification, employed stacked bidirectional LSTM layers for feature extraction which is an intuitive choice given a recurrent model’s capability of capturing global sequence-order information [28]. However, the recurrent connections and the gates also introduce a large number of parameters that need to be tuned. This can lead to overfitting since the number of training instances is limited. Moreover, since only 15 N-terminus amino acids have been considered for feature extraction, LSTM’s long-range sequence-order-effect detection capabilities remain underutilized while the parameter overhead persists [28] In this study, we do not add any recurrent layer on top of the output feature maps from the final convolutional layers to avoid this issue.

Furthermore, it is to be noted that the kernels in the final layer of each convolutional group have an effective receptive field of length 6 due to hierarchical relationship between the stacked layers (length 4 kernels to length 3 kernels) [41]. This effective receptive field should provide sufficient coverage for extracting both short-range and long-range patterns from sub-sequences of length 15. In addition, since we extract features from short sub-sequences, reducing the temporal dimension of the intermediate feature maps (outputs of the first and second convolutional layers of each group) is not required for learning higher order features. Hence, we do not add any pooling layers between the feature extraction layers within the convolutional groups [41]. The absence of pooling layers also reduces potential loss of sequence order information that can be exploited by the kernels in the final convolutional layers in the groups for detecting long-range patterns [41].

To analyse the contribution of features extracted from each of the information sources, we carry out experiments using all possible combinations of the three representations. This results in seven models (^3^*C*_1_ +^3^*C*_2_ +^3^*C*_3_) with 1, 2 or 3 convolutional groups. All these combinations are summarized in **Table 1**. The performance for our architecture using these seven combinations is reported in the following section.

**Table 1:**
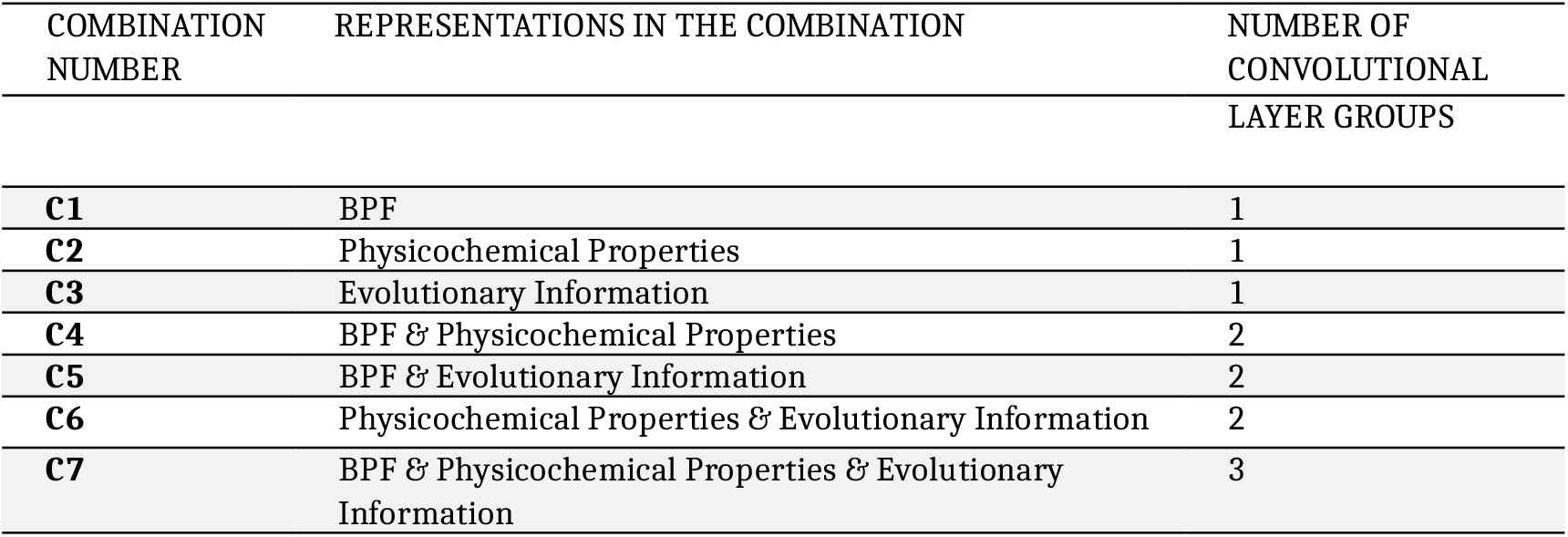
Summary of seven combinations of the three sequence representations explored in this study. On the First column of the table, we present the name of the combination, on the second column we present the name of the representations used to build the given combination, and in the third column we present the number of convolutional groups for the given combination.

For ACP-740 and ACP-240, our model’s hyperparameters are tuned on ACP-740 through cross-validation, and the same model configuration is used for ACP-240. For ACP-500 and ACP-164, hyperparameter tuning is performed on ACP-500 through cross-validation. ACP-240 and ACP-164 have been kept untouched during hyperparameter tuning to avoid performance overestimation. **Table 2** shows detailed hyperparameter configurations for different ACP identification datasets used in this study.

**Table 2:**
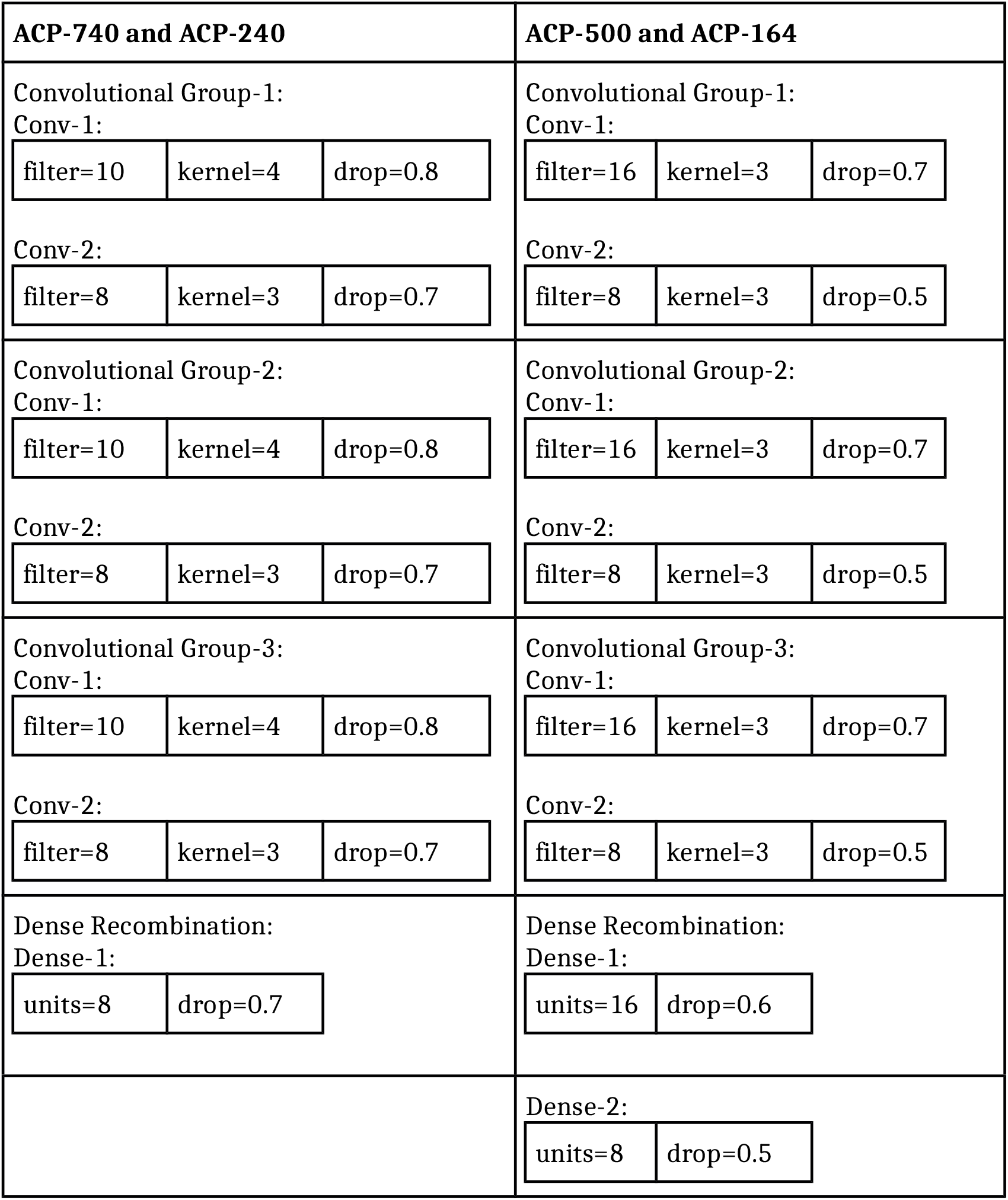
Hyperparameter configurations employed for different ACP datasets. In this table, ‘Conv’ = a convolutional layer, ‘Dense’ = a fully connected layer, ‘filter’ = number of filters in a convolutional layer, ‘kernel’ = size of filters in a convolutional layer, ‘drop’ = dropout rate, and ‘units’ = number of neurons in a fully connected layer.

## 3. Results and Discussion

In this section, we present how we carry out the performance evaluation of our proposed model, our achieved results, and then discuss them.

### 3.1 Evaluation Metrics

The evaluation metrics that have been used for measuring the performance of our classification method are Accuracy, Sensitivity, Specificity, Precision, and Matthews correlation coefficient (MCC). These metrics are described through the following equations:

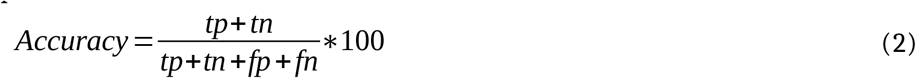

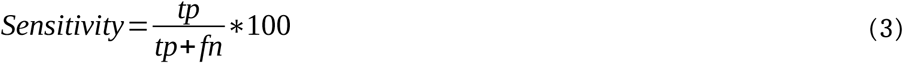

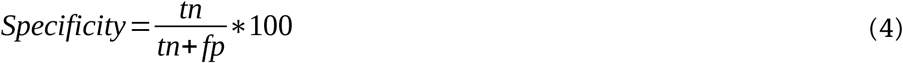

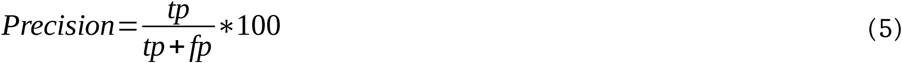

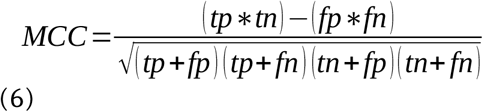

Where, *tp* is the number of correctly predicted positive instances, *tn* is the number of correctly predicted negative instances, *fp* is the number of incorrectly predicted negative instances, and *fn* is the number of incorrectly predicted positive instances. The range of values for Accuracy, Sensitivity, Specificity, and Precision is 0 to 100 percent. 100% represents an ideal classifier (totally accurate) and 0% represents the worst possible model (totally inaccurate). In addition to that, MCC has a range from −1 to +1. A value of 0 in MCC represent a random classifier with no correlation, +1 represent perfect positive correlation and −1 represents perfect negative correlation.

### 3.2 Contribution Analysis for Different Sequence Representations

For each of the representation combinations summarized in **Table 1**, we have performed experiments on ACP-740 and ACP-240 using 5-fold-cross validation, and the corresponding results are reported in **Table 3** and **Table 4**, respectively. For ACP-500 and ACP-164, we train and tune the models on ACP-500 and test them on ACP-164. The corresponding results are reported in **Table 5**.

**Table 3:** Results achieved using 5–fold cross validation for ACP-740 dataset.

| Combination | Accuracy | Sensitivity | Specificity | Precision | MCC |
| --- | --- | --- | --- | --- | --- |
| C1 | 76.0 | 78.9 | 73.0 | 75.0 | 0.52 |
| C2 | 73.1 | 74.7 | 71.3 | 72.8 | 0.46 |
| C3 | 81.1 | 81.3 | 80.7 | 81.3 | 0.62 |
| C4 | 76.9 | 75.7 | 78.4 | 78.2 | 0.54 |
| C5 | 84.0 | 87.6 | 80.3 | 82.0 | 0.68 |
| C6 | 81.8 | 82.9 | 81.1 | 81.8 | 0.64 |
| **C7** | **86.0** | **88.9** | **83.1** | **84.4** | **0.72** |

**Table 4:**
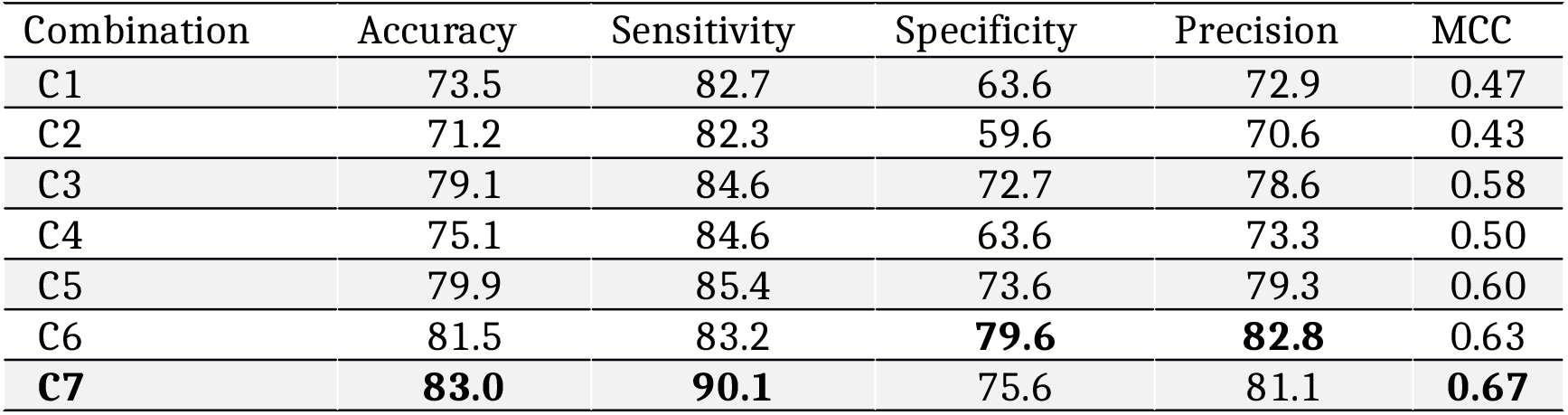
Results achieved using 5–fold cross validation for ACP-240.

| Combination | Accuracy | Sensitivity | Specificity | Precision | MCC |
| --- | --- | --- | --- | --- | --- |
| C1 | 73.5 | 82.7 | 63.6 | 72.9 | 0.47 |
| C2 | 71.2 | 82.3 | 59.6 | 70.6 | 0.43 |
| C3 | 79.1 | 84.6 | 72.7 | 78.6 | 0.58 |
| C4 | 75.1 | 84.6 | 63.6 | 73.3 | 0.50 |
| C5 | 79.9 | 85.4 | 73.6 | 79.3 | 0.60 |
| C6 | 81.5 | 83.2 | **79.6** | **82.8** | 0.63 |
| **C7** | **83.0** | **90.1** | 75.6 | 81.1 | **0.67** |

**Table 5:** Results achieved using independent test for ACP-500/164 dataset

| Classifier | Accuracy | Sensitivity | Specificity | Precision | MCC |
| --- | --- | --- | --- | --- | --- |
| C1 | 83.8 | 85.4 | 81.6 | 82.3 | 0.67 |
| C2 | 74.2 | 77.9 | 70.6 | 72.6 | 0.49 |
| C3 | 89.0 | 91.4 | 86.6 | 87.2 | 0.78 |
| C4 | 85.6 | 88.7 | 82.6 | 83.6 | 0.71 |
| C5 | 90.0 | 93.7 | 86.3 | 87.3 | 0.80 |
| C6 | 88.4 | 89.4 | **86.7** | **87.1** | 0.76 |
| **C7** | **91.0** | **97.6** | 84.2 | 86.0 | **0.82** |

As shown in **Table 3**, for the ACP-740 dataset, among the single-representation combinations (C1, C2, and C3), the representation depicting evolutionary information of the amino acid residues (C3) performs better compared to BPF and physicochemical-based representations (C1 and C2) on all six performance measures. As shown in Tables 4 and 5, similar results are observed for single representation models for ACP-240 and ACP-164. These results indicate when it comes to feature extraction from a single peptide representation, evolutionary information contributes the most for separating the ACPs from the non-ACPs compared to BPF and physicochemical-based representation.

Among the two-representation combinations (C4, C5, and C6), C5 (BPF + evolutionary) and C6 (physicochemical property + evolutionary information) performs better than C4 (BPF + physicochemical property) which further underscores the importance of the features extracted from evolutionary information for ACP identification. Moreover, C5 and C6 (two-representation combinations containing evolutionary information) perform better than C3 (the best performing single-representation combination containing evolutionary information only). This aspect of the results manifests that our proposed joint pattern extraction strategy from multiple representations through parallel-convolutional-groups can effectively embellish the features learned from a strong primary representation (evolutionary information in this case) through potential ambiguity resolution using other secondary representations (BPF and physicochemical property-based information in this case).

This hypothesis has been further corroborated by the performance of the all-representation combination (C7) on all datasets. As shown in Tables 3, 4, and 5, the model trained on C7 consisting of three parallel convolutional groups that extract features from all three representations performs better than the other combinations (C1 to C6). Therefore, we use this all-representation combination model to train ACP-MHCNN and compare its performance with state-of-the-art methods in the following subsection. To provide more insight into our achieved results, we present receiver operating characteristic (ROC) curves for our achieved results. The ROC curve for ACP-740 (using 5-fold cross validation), ACP-240 (using 5-fold cross validation), and ACP-164 (using ACP-500 as the training dataset) are shown in Figures 2, 3, and 4, respectively.

**Figure 2:**
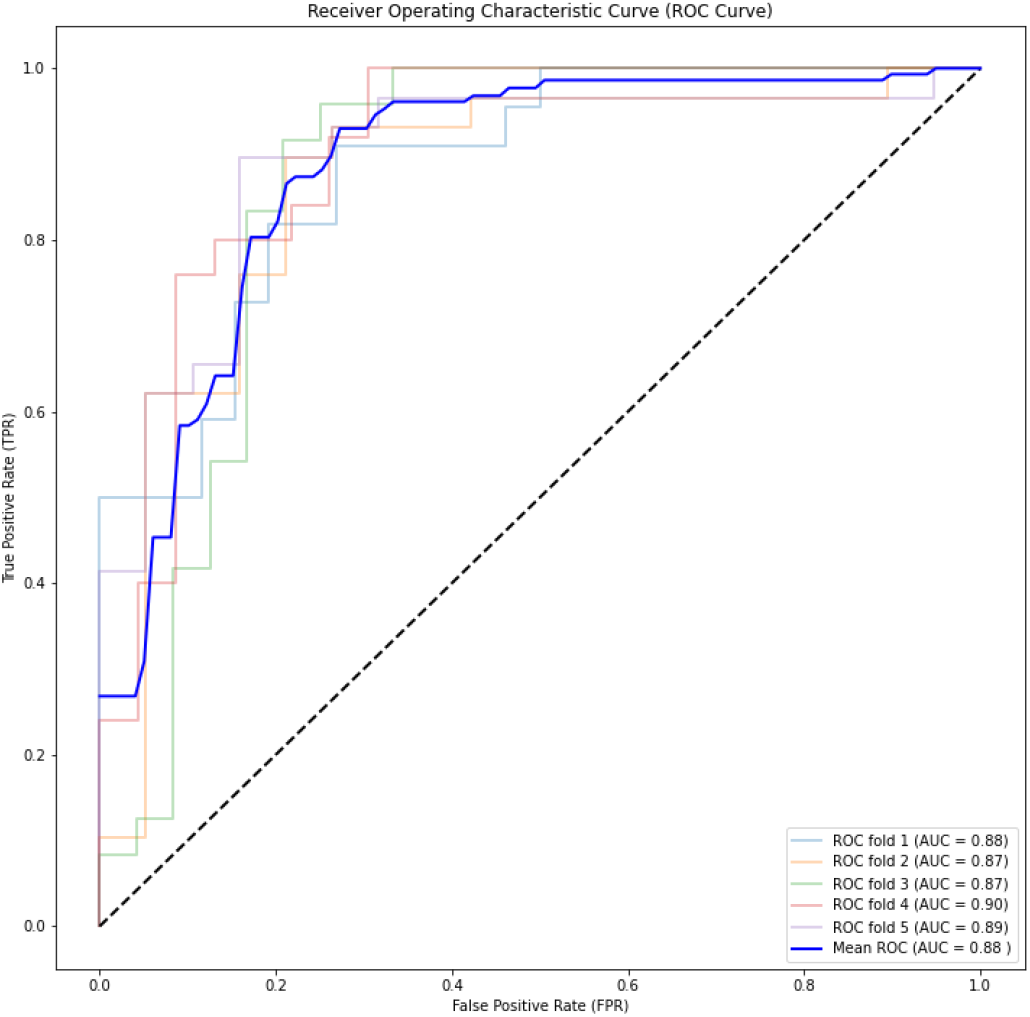
ROC curve for ACP-740 dataset for the 5-fold cross-validation on the experiment.

**Figure 3:**
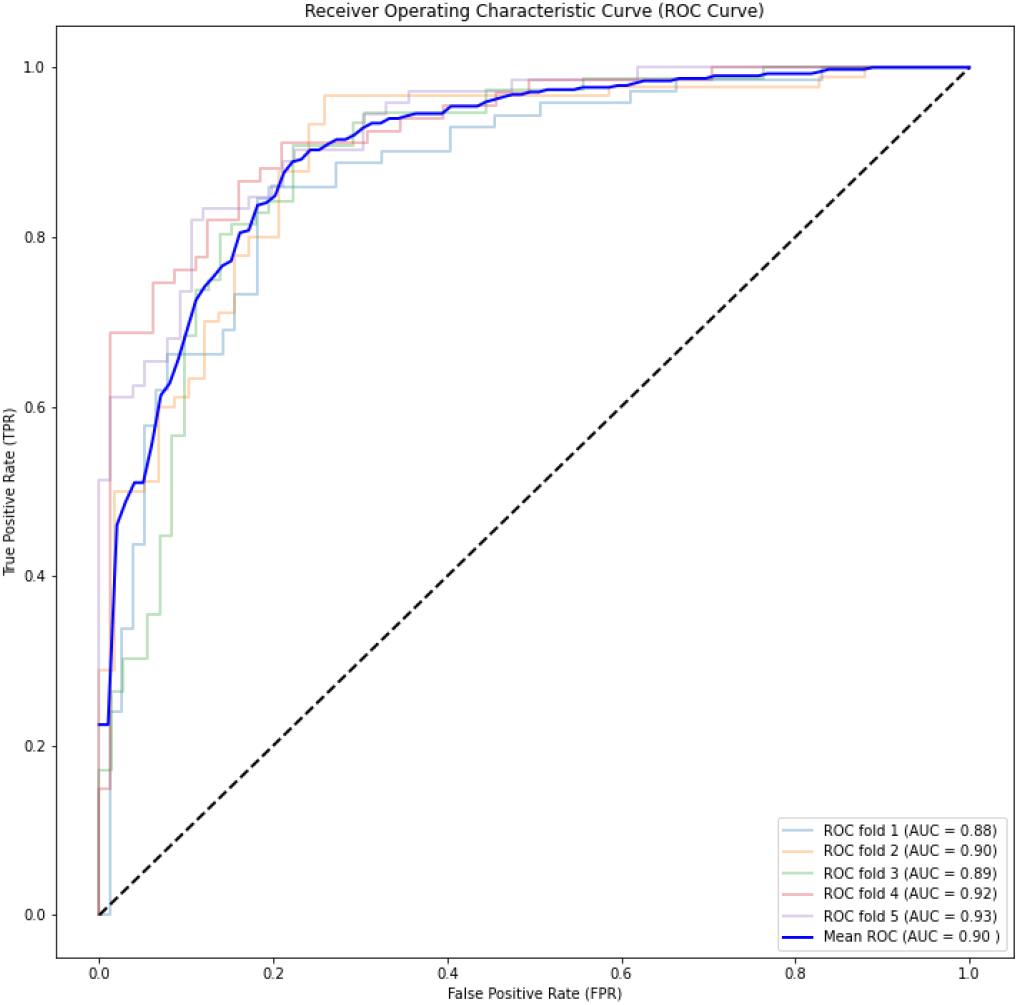
ROC curve for ACP-240 dataset for the 5-fold cross-validation on the experiment.

**Figure 4:**
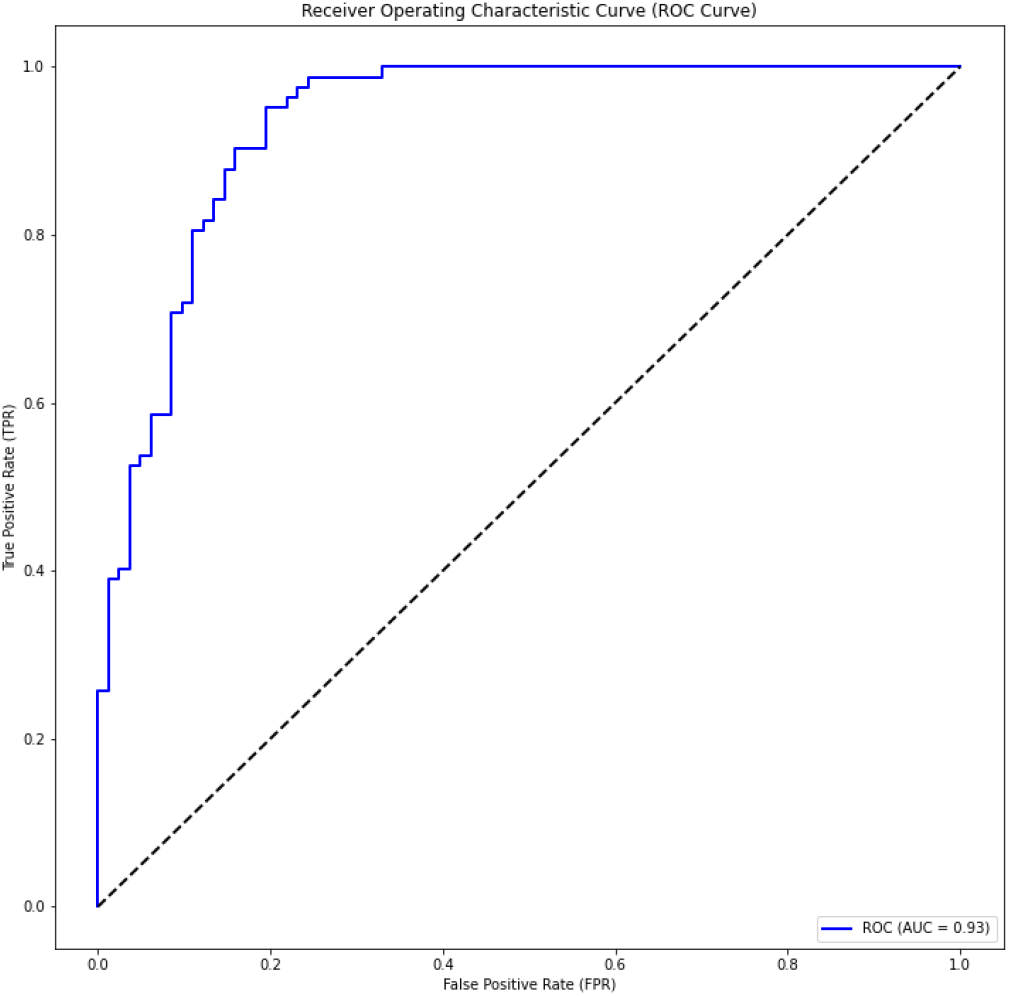
ROC curve for ACP-500/164. Here We used ACP-500 as a training dataset and ACP-164 as a testing dataset on the experiment.

As shown in these figures, we constantly achieve very high Area Under the Curve (AUC) value. We achieve 0.90, 0.88, and 0.93 for ACP-740, ACP-240, and ACP-164, respectively. The consistent AUC achieved on these three benchmarks using different evaluation methods demonstrates the generality of our proposed model. In addition, achieving 0.93 in AUC on ACP-164 which is an independent test set demonstrates the potential of ACP-MHCNN on identifying ACP for new unseen samples.

### 3.3 Comparison with State-of-the-art Methods

We compare ACP-MHCNN with ACP-DL as the state of the art and also the only DL based ACP identification model proposed to date [28]. Yi et al. tested their proposed ACP-DL on ACP-740 and ACP-240 datasets using 5-fold cross-validation. We use the same evaluation strategies and metrics for a fair comparison while estimating our ACP-MHCNN’s performance on ACP-740 and ACP-240 datasets. To investigate the generality of ACP-MHCNN even furtherACP-MHCNN, we compare it with ACP-DL on ACP500/ACP164 dataset as well. In this experiment, ACP-500 is used for training and tuning the model, and ACP-164 is used as independent dataset. During all these experiments, ACP-DL is trained using the implementation details available in the accompanying Github repository (https://github.com/haichengyi/ACP-DL).

Comparison between ACP-MHCNN and ACP-DL on all the datasets is shown in **Table 6**. As shown in this table, ACP-MHCNN outperforms ACP-DL on all datasets for every evaluation metric. To be precise, on ACP-740, ACP-MHCNN scores 6.0%, 7.5%, 4.5%, 4.7%, and 0.12 more than ACP-DL in terms of accuracy, sensitivity, specificity, precision, and MCC, respectively. Similarly, on ACP-240 ACP-MHCNN scores 1.8%, 6.0%, 4.4% and 0.02 more than ACP-DL in terms of accuracy, specificity, and MCC, respectively.

**Table 6:**
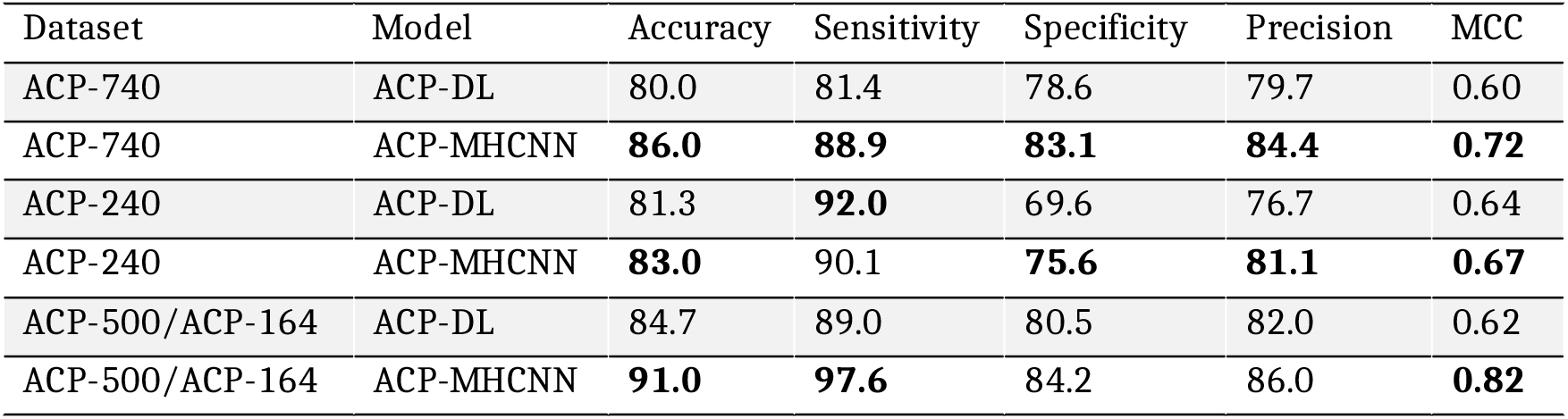
Comparing the results achieved for ACP-BNN to ACP-DL as the state-of-the-art anticancer peptide predictor.

| Dataset | Model | Accuracy | Sensitivity | Specificity | Precision | MCC |
| --- | --- | --- | --- | --- | --- | --- |
| ACP-740 | ACP-DL | 80.0 | 81.4 | 78.6 | 79.7 | 0.60 |
| ACP-740 | ACP-MHCNN | **86.0** | **88.9** | **83.1** | **84.4** | **0.72** |
| ACP-240 | ACP-DL | 81.3 | **92.0** | 69.6 | 76.7 | 0.64 |
| ACP-240 | ACP-MHCNN | **83.0** | 90.1 | **75.6** | **81.1** | **0.67** |
| ACP-500/ACP-164 | ACP-DL | 84.7 | 89.0 | 80.5 | 82.0 | 0.62 |
| ACP-500/ACP-164 | ACP-MHCNN | **91.0** | **97.6** | 84.2 | 86.0 | **0.82** |

ACP-MHCNN also significantly outperforms ACP-DL on the ACP-500/ACP-164 dataset that was used to investigate the generalizability of our model. On ACP-500/ACP-164 ACP-MHCNN outperforms ACP-DL by 6.3%, 8.6%, 3.7%, 4.0%, and 0.20 in terms of accuracy, sensitivity, specificity, precision, and MCC respectively. ACP-MHCNN and its relevant codes as well as the datasets used in this study are all publicly available at: https://github.com/mrzResearchArena/Anticancer-Peptides-CNN.

## 4. Conclusion

In this study, we propose a new deep neural network architecture called ACP-MHCNN consisting of parallel convolutional groups which jointly learn and combine features from three different peptide representation methods for accurate identification of ACPs. The architecture extracts sequence-based features from residue-order information (using BPF representation), physicochemical property-based features from 31 bit-vector representation of the residues (elements of these vectors depict various physicochemical properties of the amino acids) and evolutionary features from BLOSUM62 matrix-based representation of the peptides.

Although hand-engineered features extracted from these information sources have been successfully employed for ACP identification, automatic feature extraction has hardly been explored for this problem. Before this study, ACP-DL was the only method that has used deep feature extraction for ACP identification [28]. It has used recurrent layers for extracting features from two residue-order-based peptide representations and leaves significant room for improvement. In the current study, we attempt to address the limitations of ACP-DL by improving the sequence representation and feature extraction methods. For sequence representation, we consider the residues’ evolutionary and physicochemical characteristics alongside their ordering so that the downstream feature extraction layers can embed the sequences in spaces with additional discriminative information. For feature extraction, we jointly train three parallel convolutional layer groups so that the combined feature vector contains discriminative patterns extracted from three sources.

The positive effects of these improvements are manifested in the experimental results obtained on well-established ACP identification datasets, where ACP-MHCNN has significantly outperformed ACP-DL using different evaluation measures for every dataset investigated in this study. Hence, we believe our current study’s findings add significantly to the existing knowledge on computational method development for ACP identification. ACP-MHCNN, its relevant codes, and the datasets used in this study are all publicly available at: https://github.com/mrzResearchArena/Anticancer-Peptides-CNN.

## Author Contributions

S. Ahmed conceived and initiated this study. S. Ahmed and R. Muhammod performed the experiments. S. Ahmed, S. Adilina and A. Dehzangi wrote the manuscript. Zahid Hossain helped with literature review. A. Dehzangi, S. Shatabda mentored and analytically reviewed the paper. All the authors reviewed the article.

